# Roles for Phosphatase PP4 in Rhythmicity and Compensation in the Neurospora Circadian System

**DOI:** 10.64898/2026.05.23.726942

**Authors:** Adrienne K. Mehalow, Ziyan Wang, Scott A. Gerber, Jennifer J. Loros, Jay C. Dunlap

## Abstract

The circadian clock is a highly conserved timer which allows organisms to anticipate future conditions driven by the 24-hour day on planet Earth. Circadian clocks from fungi to mammals are based on a transcription-translation feedback loop (TTFL) molecular architecture. Progressive phosphorylation of core clock proteins can alter both their activity and stability and is required for all known circadian TTFLs. The mechanism for kinase control of circadian period has been extensively studied; however, the mechanism(s) whereby phosphatases alter period remain less studied. Based on the observation that strains of *Neurospora crassa* lacking phosphatase *pp*4 display a short circadian period, we investigated regulation of *pp*4 and its role in the clockworks. In addition to period shortening, loss of *pp*4 results in a significant loss of both temperature and nutritional compensation, consistent with substrates within the core clock. We identify the clock-relevant PP4 phosphatase holoenzyme as a heterotrimer, and identify two activators of PP4 which regulate circadian period. Biochemical and cell biological analyses suggest that PP4 acts in the nucleus, and a mass spectrometry-based screen identified NCU07414, a member of the HSP40 chaperone system, as a novel PP4 binding partner whose loss results in a dramatically shortened period of the clock when ablated. A model consistent with the data suggests that PP4 may act in opposition to kinases to influence the rate of accumulation of the clock-relevant phosphorylations that determine circadian period.

## INTRODUCTION

The circadian clock facilitates organization and entrainment of rhythmic biological processes with Earth’s daily environmental changes [1]. Timekeeping provides such a significant fitness advantage that circadian oscillators have appeared independently at least three times during evolution [2]. In humans, misalignment of the circadian clock and daily activity is a significant health risk that is associated with metabolic syndrome, psychiatric disease, and the development of cancer [3–5]. Eukaryotes from fungi to mammals all share a circadian clock structure based on a similar regulatory architecture [6]. In this mechanism, positive-acting heterodimeric transcription factors drive expression of target mRNAs coding for negative-acting proteins; these form complexes that include casein kinase 1 and act via phosphorylation to feedback and suppress the activity of the original positive elements. The result is a regular oscillation of gene transcription that is tuned to occur once per 24-hour day.

The core clock is a transcription and translation negative feedback loop (TTFL) [7]. In Neurospora the positive acting heterodimeric transcription factor WCC (composed of the WC-1 and WC-2 proteins) is bound to target promoters in the late night/early morning where it drives production of transcripts. One of the target genes is *frequency (frq)* which encodes a scaffold for formation of the negative arm complex of the clock. *frq* transcripts peak in abundance in the mid-day, with FRQ protein peaking ~4-6 hours later. FRQ complexes with the FRQ-Interacting RNA Helicase (FRH), which acts as a nanny protein to stabilize FRQ, and translocates into the nucleus where it forms the negative arm FFC complex by associating with casein kinase 1a (CK-1a) [8]. FRQ becomes progressively phosphorylated and in the mid to late day the FFC brings CK1 to WCC in the nucleus. During the night WCC binding to DNA is weakened due to phosphorylation and it is released from DNA. WCC is further exported from the nucleus where it can be dephosphorylated in preparation for the next cycle. Meanwhile FRQ becomes inactivated by hyperphosphorylation, which relieves repression, and it is eventually turned over although this turnover is not an obligate step in the cycle [9]. The next cycle begins just before dawn with formation of the WC-1/WC-2 heterodimer which is imported to the nucleus.

Along with persistence of circa 24 hour rhythmicity under constant environmental conditions and the ability to be entrained by environmental changes, one of the defining properties of a circadian clock is its ability to maintain nearly the same period length over a physiologically relevant range of temperatures or nutrition, characteristics known as temperature compensation (TC) or nutritional compensation (NC) [10–12]. A mechanistic explanation for compensation remains lacking even after decades of research in chronobiology, perhaps because the phenomenon is complex and has been imperfectly defined. For instance, some but not all alleles of *frq* alter TC [13]; however, a typical garden variety temperature sensitive allele of a clock protein might present as a TC-mutant if its clock-relevant activity changes as a function of temperature. Early conceptual work on compensation posited that TC and NC might both be reflections of a single basic compensation mechanism thereby conflating the two phenomena. However, a recent systematic screen for genes involved in NC revealed that this phenomenon is genetically separable from TC; that is, not every gene that affects NC also affects TC and vice versa [8]. Aside from core clock proteins, mutations shown to affect NC have implicated the transcription factor CSP-1 and co-factor RCO-1, the RNA helicase PRD-1, and proteins involved in alternative splicing and polyadenylations such the CFIm complex [8], but no kinases are associated with NC. In contrast, and again excluding core clock proteins that are likely to be targets of TC, *Neurospora* genes known to function in TC include only the kinases CK1, CK2, and Chk2 [12]. These facts suggest an ultimate mechanism for TC is likely to end with a protein(s) of the core clock (e.g. in the FCC or WCC) which is being altered by one or more phosphorylation events. Perhaps significantly, no phosphatases have been implicated in NC or TC.

PP4 is part of the PPP family of multimeric phosphatases which are conserved from yeast to humans [14, 15]. All PPP phosphatases share a highly conserved catalytic domain and function as a complex of a single catalytic subunit associated with one or more regulatory subunits. For the sake of clarity, we will refer to the multimeric phosphatase as PP4 holoenzyme, and the individual subunits using their gene/protein nomenclature for the respective organism (*pp*4/PP4, *reg*3/REG3, etc.). *Neurospora pp*4 deletion strains (knockouts) are viable and result in short circadian period, defective clock output, reduced growth, and defects in sexual reproduction [16, 17]. In contrast, PP4 is an essential phosphatase in mammals where it functions in double-strand break repair and telomere maintenance, and in Drosophila where it is required for centrosome maturation [18]. At least five regulatory subunits are known in humans (PP4R1, PP4R2, PP4R3A, PP4R3B, PP4R4), while in yeast there are two regulatory subunits (Psy2p, Psy4p) [19]. The combinatorial of regulatory subunits in the active phosphatase defines substrate specificity, with the possibility that multiple active phosphatase holoenzymes may exist simultaneously [20]. Generation of the active PP4 holoenzyme requires latency chaperones, activating proteins, PTMs, and assembly of the catalytic subunit with appropriate regulatory subunits [21]. Briefly, in mammals nascently synthesized PP4 catalytic subunits are stabilized by interactions with alpha4, which acts as a chaperone to prevent unregulated activity. Subsequent complex assembly and activation steps are performed by TIPRL and Ypa1/2. PPP catalytic subunits, including PP4, share a C-terminal DYFL amino acid motif which is often subject to methylation on the terminal leucine residue by a phosphate methyl esterase. Addition of this methyl group is thought to be required for assembly of the active phosphatase [22]. Beyond assembly and activation, the mechanism by which PP4 identifies targets for dephosphorylation is broadly unknown. Recent work in mammals suggests that substrates presenting the Short Linear Motifs (SLiMs) FxxP and MxxP interact with regulatory subunits PPP4R3A and PPPR3B to confer selection [23].

In this study we pursued an unexpected alteration in the circadian period previously described for a *pp*4 knockout [17, 24]. Surprisingly, nutritional increases lead to period shortening and using this knowledge we observed a significant TC defect that was masked in prior screens. Regulatory subunits REG2 (NCU00900) and REG3 (NCU06389) that are part of the PP4 holoenzyme were identified. The PP4 holoenzyme is regulated by upstream activators STK-2 (NCU04810) and TIPA (NCU00109), but not by leucine C-terminal methylation of the catalytic subunit. Localization of WC-1 and WC-2 is not altered *in vivo* in a *pp*4 knockout, challenging a prior model in which WCC nuclear entry was regulated by PP4 [17], and fluorescent tagging of REG3 confirmed an exclusively nuclear localization. Because homologues of REG3 are responsible for selection of phosphatase substrates, we propose the activity of the PP4 holoenzyme is nuclear and the clock-relevant PP4 substrate in *Neurospora* will also be found in this compartment. Although PP4 interacting partners are notoriously difficult to identify [25], we used epitope-tagged REG3 in an immunoprecipitation/mass spectrometry-based screen to identify potential phosphatase-substrate complexes. Genetic screening of the candidate interactors identified *djc*-4 (NCU07414), a Heat Shock Protein 40 (HSP40) protein, whose knockout displays an extremely short period of ~12-14 hours, to our knowledge the shortest period length associated with a single gene knockout to date.

## RESULTS

### Loss of *pp*4 impacts circadian period length as well as circadian temperature and nutritional compensation

A recently completed a screen of existing and newly annotated Neurospora protein phosphatases for involvement in circadian period length and temperature compensation [26] uncovered a strikingly short period of 17.3 +/−0.3 hours at 25°C for Δ*pp*4 (Fig 1A) and partial loss of temperature compensation leading to substantial overcompensation such that the period lengthened with increasing temperature. (Prior literature has made a hash of nomenclature. NCU08301 (FGSC12453/FGSC12454) encodes a calcineurin-like protein phosphatase called protein phosphatase X in Borkovich *et al*., 2004 [27], and encoded by *pp*4 in Cha *et al.*, 2008 [17] and by *pp*h-4 in Ghosh *et al.*, 2014 [16]. NCU08301 is called *pp*4 in FungiDB and the Saccharomyces ortholog is *pp*h3. As Cha *et al.*, 2008 have historical priority for the gene name we use *pp*4 for the gene and PP4 for the protein.) While a short period is consistent with prior estimates based of knockouts of *pp*4 in *Neurospora* [17, 24] and analysis of PP4 in other model organisms [28], our estimate was much shorter than reported by prior work (~18 hrs in *Neurospora*) suggesting additional factor(s) were affecting period length.

**Figure 1.**
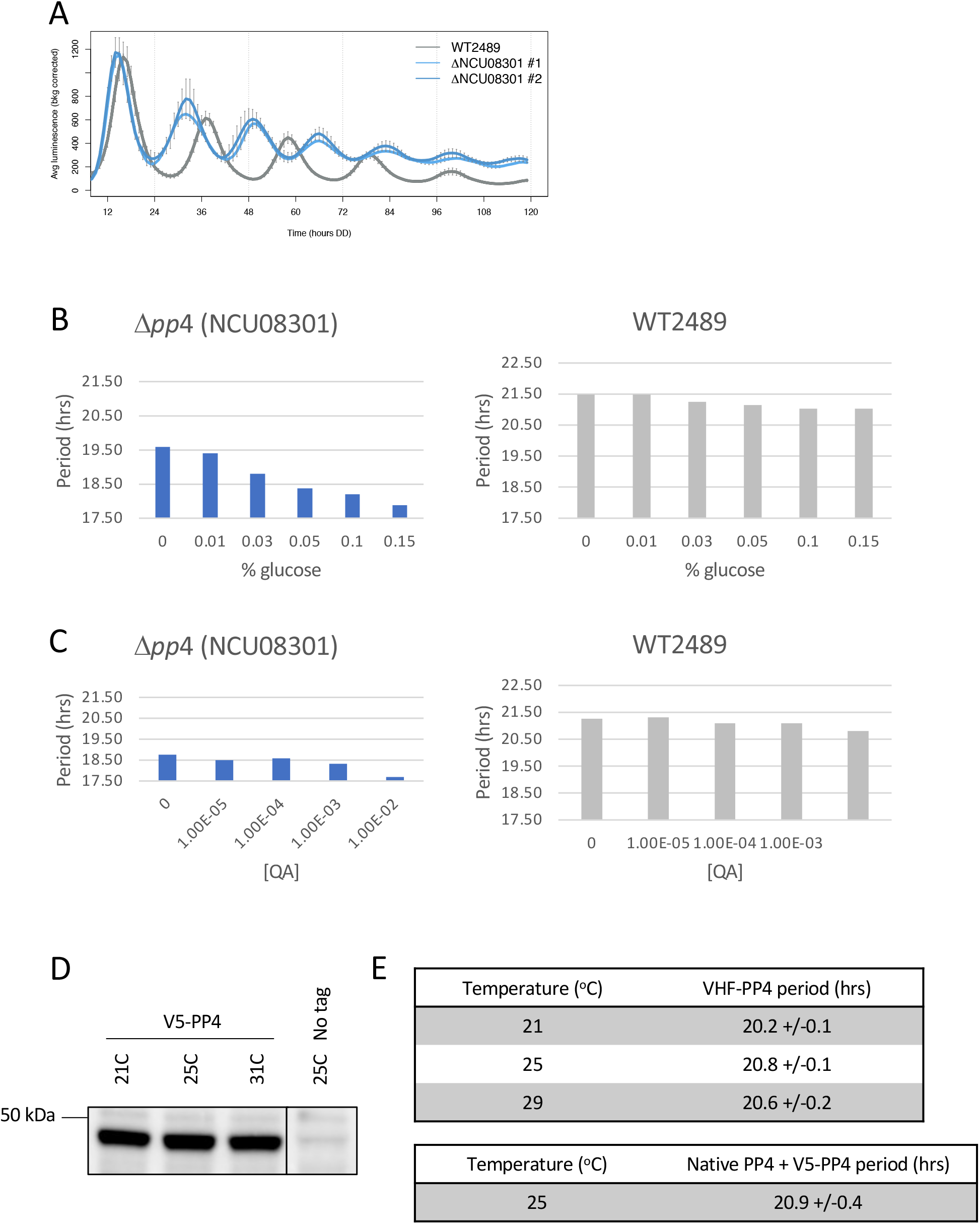
*pp*4 deletion strains exhibit defective nutritional compensation. (A) Loss of *pp*4 (NCU08301) produces short period of 17.1 +/−0.3 hrs at 25C, compared with wildtype control period of 21.1 +/−0.1 hrs. (B) The short period in *pp*4 deletions is modified by nutrition. Increasing percentage of glucose from 0% to 0.15% shortens period by greater than 1.5hrs in the knockout, while period of wildtype controls was altered by less than 0.5 hrs. (C) Increasing percentage of quinic acid (QA) from 0M to 0.01M shortened period in *pp*4 deletions. This effect was less pronounced than glucose, with period shortening of approximately 1 hr. Wildtype controls were altered by 0.5 hrs under the same conditions. (D) V5 N-terminal tagged PP4 is expressed at levels that do not change appreciably with temperature. Anti-V5 Western blot of V5-PP4 targeted to the native locus is shown. Results were similar for VHF-PP4 tagging. Band size is consistent with a predicted size of 46 kDa for PP4. (E) Appending an N-terminal tag has minimal impact on PP4 activity across temperatures. Period of VHF-PP4 was assessed by tracking cbox-luc reporter activity at 25°C under constant darkness. Periods for VHF tagged strains are shown; V5-PP4 tagging resulted in similar period lengths. Insertion of a second copy of V5-tagged PP4 at the *csr*-1 locus does not appreciably shorten period.

The reporter for our circadian period screen uses the *frq* clock-box (cbox) promoter to drive luciferase expression to report WCC activity in live *Neurospora* cultures growing on solid media [29]. Different media preparations have been used for growth in this reporter system although none have been found to significantly alter wildtype period. Media for our screen [26] contained both 0.01M quinic acid (QA) and 0.01% glucose and yielded a wildtype period of 21.3 +/−0.1 hours at 25°C, similar to the 21.74 +/− 0.29 hour period reported for media containing only glucose [24]. Noting this difference, we tested a range of glucose and QA concentrations at 25°C. Surprisingly the period in Δ*pp*4 became shorter as either glucose or QA levels increased, suggesting that nutritional compensation is impaired in the absence of the phosphatase (Fig 1B and 1C).

We generated several constructs with N-terminally tagged PP4 at the native locus in order to interrogate protein interactions. The abundance of V5-tagged PP4 was not altered by temperature between 21°C and 31°C (Fig 1D). Circadian period of either the V5 or V5-His6-3X FLAG tagged PP4 was similar to wildtype controls (Fig 1E). A modest period lengthening effect of overexpression of PP4 on has been reported for human cells in culture [28]. To determine if Neurospora is affected by increased PP4 expression, we inserted V5-tagged PP4 at the *csr*-1 locus of an otherwise wildtype strain expressing the cbox-luc reporter at *his*-3. Even with twice the gene dosage of *pp*4, period remained similar to wildtype (Fig 1E).

### Absence of PP4 does not alter WCC localization *in vivo*

PP4 is proposed to dephosphorylate WCC to allow nuclear re-entry of the transcription factor [17]. Our lab recently developed fluorescently tagged WC-1 and WC-2 proteins for *in vivo* imaging of positive arm dynamics [30]. We crossed the *pp*4 deletion strain with a WC-1 strain bearing a C-terminal mCherry tag, and a WC-2 strain bearing a C-terminal mApple tag and imaged the growing hyphal tips. At 25°C WC-1 and WC-2 localization was predominantly nuclear, and there was no difference in the localization of either protein when PP4 was absent (Figure 2A, 2B). Due to the challenges of *in vivo* imaging in Neurospora, we are unable to determine with statistical certainty if total abundance of WC proteins is unchanged; however, if differences are present they are not large.

**Figure 2.**
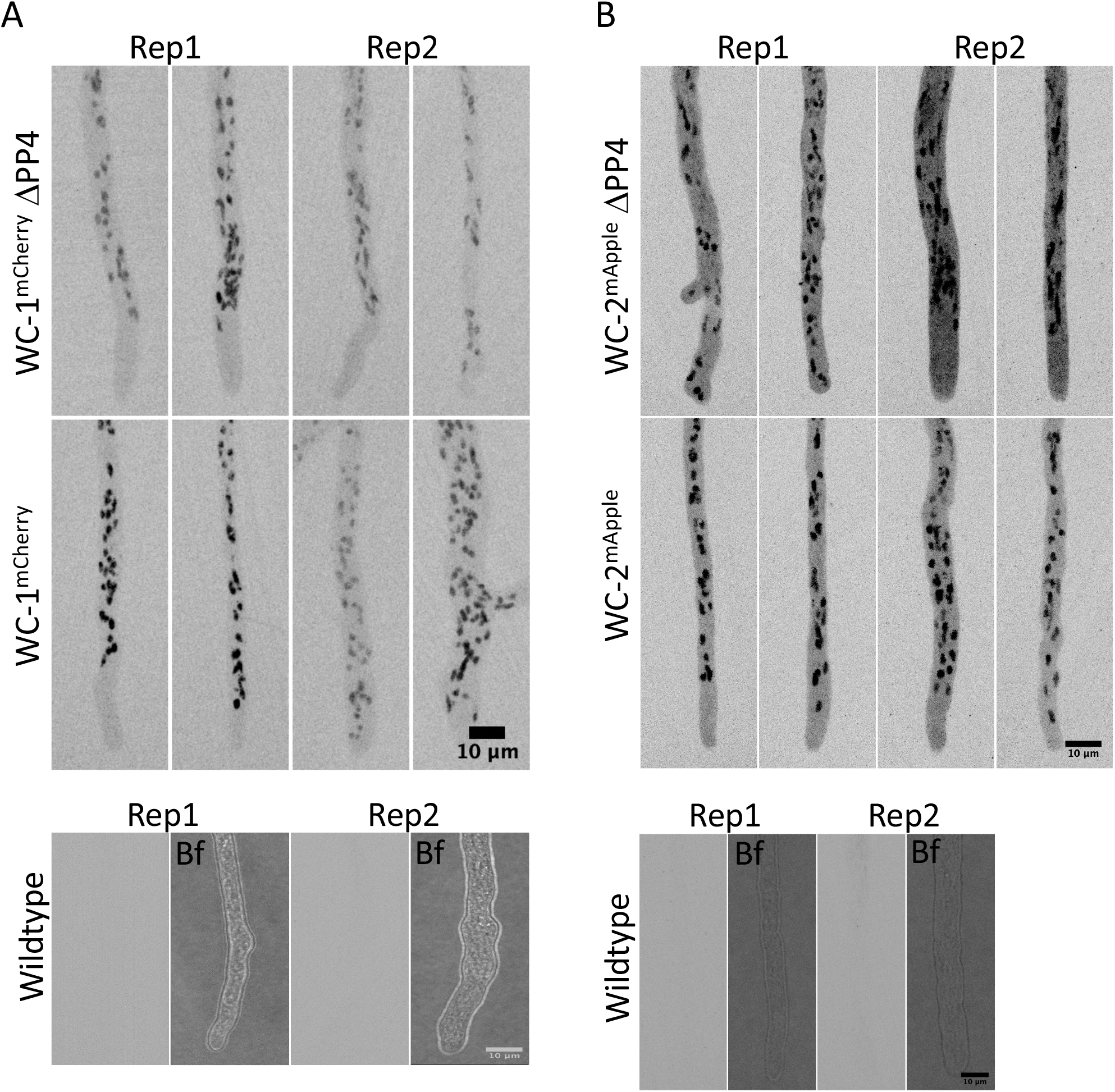
Localization of WC-1 and WC-2 is not affected by loss of *pp*4. (A) WC-1 C-terminally tagged with mCherry localizes to the nucleus in vivo at 25°C, as previously reported. Deletion of *pp*4 does not alter this localization. (B) WC-2 C-terminally tagged with mApple also localizes to the nucleus at 25°C; deletion of *pp*4 has no effect on localization. For (A) and (B) representative images of hyphal tips from two unique progeny growing on solid media are shown for each genotype. Images were obtained with 561nm excitation and are presented at maximum Z-projections of 35 slices at 0.3um intervals. An untagged sibling with normal PP4 expression is shown for control.

### Mass spectroscopy and data mining identified PP4 holoenzyme subunits

Multiple regulatory subunits for PP4 have been identified in yeast, mammals, and Drosophila, but orthologous proteins have not been assigned in *Neurospora*. However, a recent liquid chromatography-tandem mass spectrometry (LC-MS/MS) based screen used the nonselective PPP inhibitor microcystin-LR (MCLR) to capture phosphatase catalytic subunits and their associated proteins from several organisms including human tissue and of *Neurospora crassa* lysates [31]. We hypothesized that PP4 and its regulatory subunits would be included in this data set because of structural similarity shared among PPP catalytic subunits. Of the 27 unique *Neurospora* proteins identified by LC-MS/MS, 8 were annotated PPP family catalytic subunits and PP4 phosphatase was indeed present. Of the remaining proteins, 3 were annotated as PP2A-associated proteins, 5 had functions not assigned to phosphatase regulation, and 11 were uncharacterized or assigned “domain of unknown function”. Relying on OrthoMCL homology assignment (Chen et al. 2006 as implemented at FungiDb [32] (www.fungibd.org)) we selected candidate PP4 subunits and identified NCU00900 and NCU06389 as the most likely to be PP4 regulatory subunits. By homology we assign NCU00900 as *reg*2 encoding the scaffolding subunit, and NCU06389 as *reg*3 encoding the subunit involved in substrate recognition. Knockouts of regulatory subunits necessary for PP4 holoenzyme function should phenocopy Δ*pp*4, and indeed we found that loss of NCU00900 (*reg*2) or NCU06389 (*reg*3) produced a short period identical to loss of the PP4 catalytic subunit (Fig3A, 3C); double knockouts of either *reg*2/*pp*4 or *reg*3/*pp*4 failed to produce further shortening of the period at 25C (Fig. 3B, 3D) suggesting that a functional phosphatase complex consisting of PP4/REG2/REG3 is required for normal period and TC of the circadian clock.

**Figure 3.**
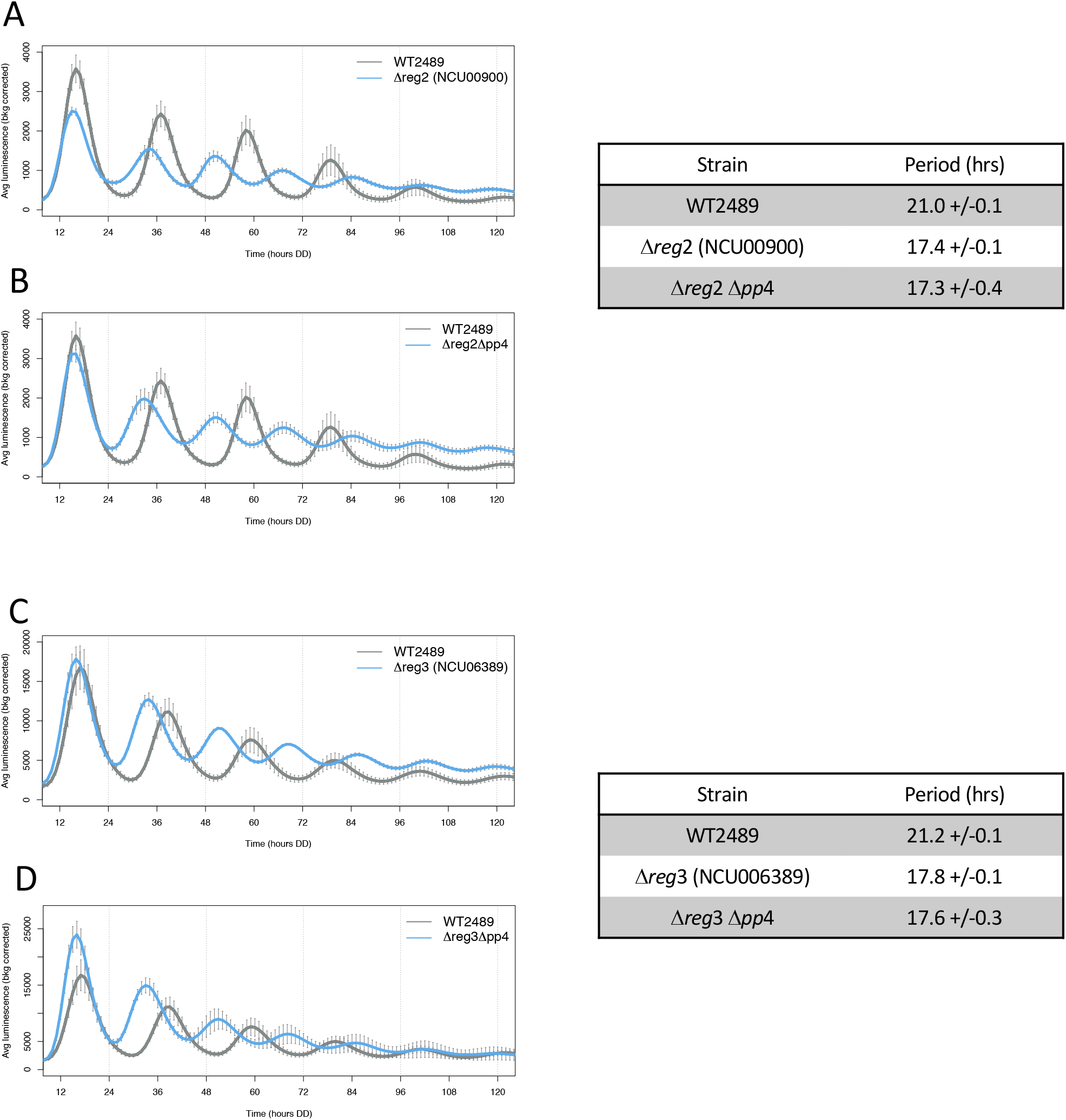
Loss of *reg*2 (NCU00900) or *reg*3 (NCU06389) replicates the short period observed in *pp*4 deletion strains. (A-D) cbox-luc luciferase reporter activity for the indicated strains at 25°C. Representative trace of one individual shown (blue) with WT2489 (gray) comparison from the same run. (A and B) Deletion of NCU00900 results in short period similar to the *pp*4 knockout strain. Double deletion of NCU00900 and *pp*4 does not further shorten period. (C and D) Deletion of NCU06389 results in short period similar to the *pp*4 deletion strain. Double deletion of NCU06389 and *pp*4 does not further shorten period.

### PP4 holoenzyme localization suggests action in the nucleus

PP4 regulatory subunit(s) should appear in the same subcellular location as the clock-relevant targets to which they bind. Further, the regulatory subunit and substrate complex must appear in the same location as the PP4 catalytic subunit at the time of substrate dephosphorylation. Because REG3 is believed to provide substrate selection for PP4, we N-terminally tagged NCU06389/REG3 with mNeonGreen at the native locus to track this protein *in vivo*. REG3^mNeonGreen^ localization was nuclear at both 20°C and 30°C (Fig. 3D), consistent with reported nuclear functions of PP4 in other organisms [33, 34]. The mNeonGreen tag did not substantially alter protein abundance with temperature (Fig. 4B). Addition of N-terminal tags including V5-His6-3X FLAG to REG3 shortened period by about one hour (Fig. 4C), indicating a minor impact on phosphatase function. However, this is distinctly different from the period of the REG3 deletion, therefore we conclude the phosphatase retains relatively normal function. Attempts to append a fluor to the N-terminus of PP4 resulted in a strain with short period, suggesting presence of a large tag interfered with function (data not shown). Taken together, live cell *in vivo* localization of WCC and REG3 are consistent with a model where the PP4 holoenzyme does not regulate the localization of WCC and acts in the nucleus.

**Figure 4.**
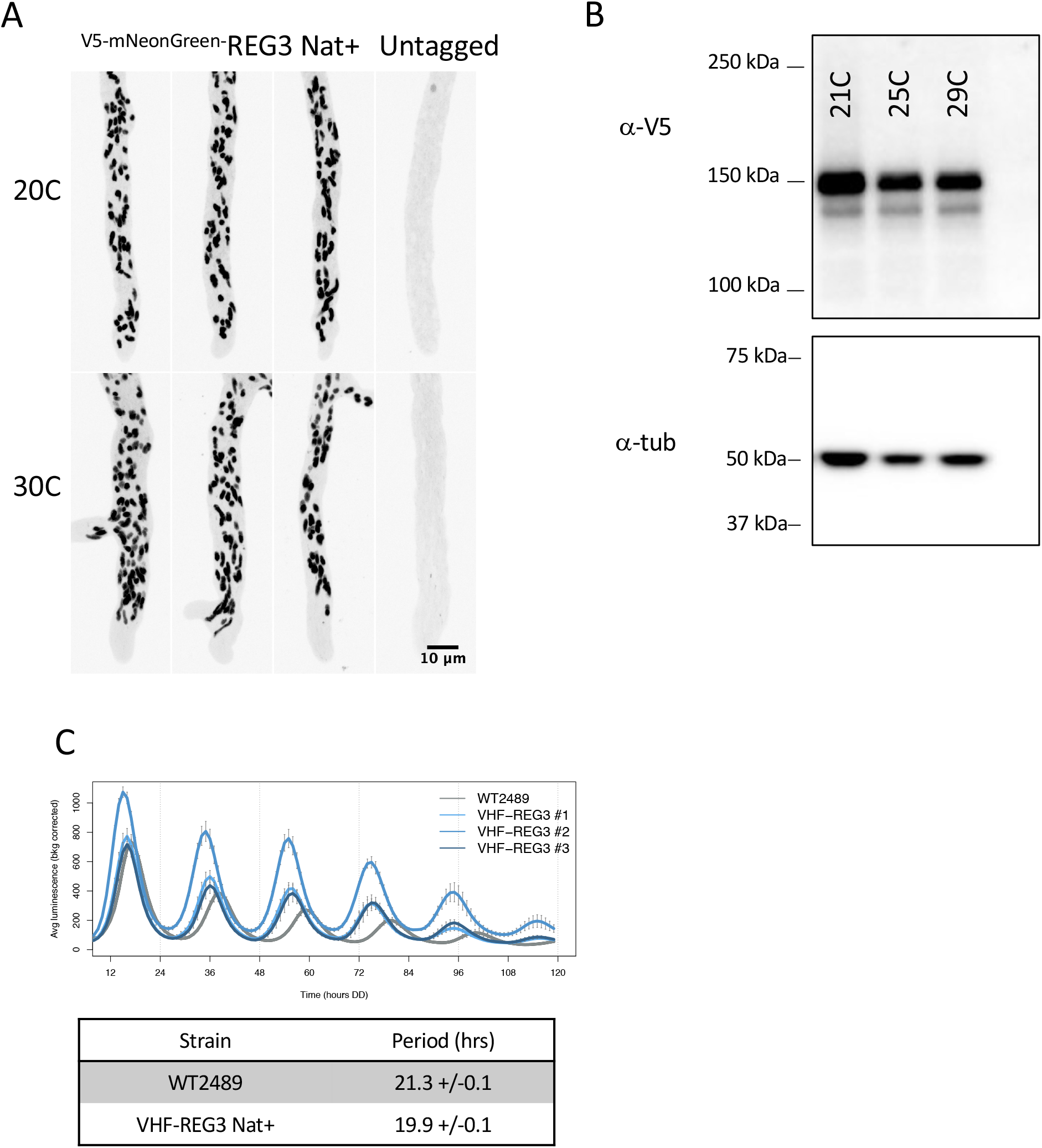
REG3 (NCU06389) is localized to the nucleus in vivo. (A) Growing hyphal tips from V5-mNeonGreen tagged REG3 strains were imaged with 488nm excitation. Images are maximum z-projections of 51 slices with 0.2um intervals. Three representative images from tagged strains and one untagged control are shown at each temperature. Nuclear localization was not altered between 20°C and 30°C growth. (B) Anti-V5 Western blot of V5-mNeonGreen REG3 strain grown at three temperatures. REG3 is expressed and the level is not appreciably changed across the temperature range tested. Anti-tubulin Western blot shown for loading control. (C) Appending an N-terminal tag to REG3 shortens period by ~1.5hrs at 25°C, as determined by cbox-luc reporter assay although period remains much longer than observed in *pp*4 deletion strains.

### The PP4 catalytic subunit is post-transcriptionally regulated in ways similar to but not identical to orthologs in other systems

PP2A-family phosphatases are post-transcriptionally regulated through many steps that appear conserved from yeast to mammals [35], and because Neurospora has orthologs to many of these components, we systematically examined them for clock effects. Newly synthesized phosphatase catalytic subunits are stabilized by interactions with phosphorylated alpha4, encoded by an essential gene which acts as a chaperone to prevent unregulated activity and premature PP4 degradation. Subsequent complex assembly and activation steps are performed by TIPRL (*Neurospora TipA*), the peptidyl-prolyl cis/trans-isomerases RRD1/2 (*Neurospora stk-2/stk-3*), and C-terminal leucine methyl transferases [22, 36]. Knockouts of *TipA* (NCU00109) are viable and displayed wild type period (Table 1). Knockouts of the paralogs RRD1 (*stks*-2, NCU04810) and RRD2 (*stk*-3, NCU03269) were examined and interestingly only Δ*stk*-2 phenocopied Δ*pp*4. To examine role(s) for the C-terminal leucine methylations that commonly regulate PP2-family phosphatases we further examined the *Neurospora* deletions of the two annotated leucine-methyl transferase enzymes, *lcm*-1 (NCU07993) and *lcm-2* (NCU5392), and one phosphate methylesterase, *pme*-1 (NCU00954), that opposes leucine methyltransferases. Knockouts of all three genes displayed wild-type circadian periods (Table 1). Taken together, these data indicate divergence of function between Neurospora and yeast/animals in that the peptidyl-prolyl cis/trans-isomerase activity of only STK-2 is required to activate the PP4, holoenzyme but once activated, neither leucine methylation nor demethylation is required for PP4 phosphatase activity.

**Table 1:**
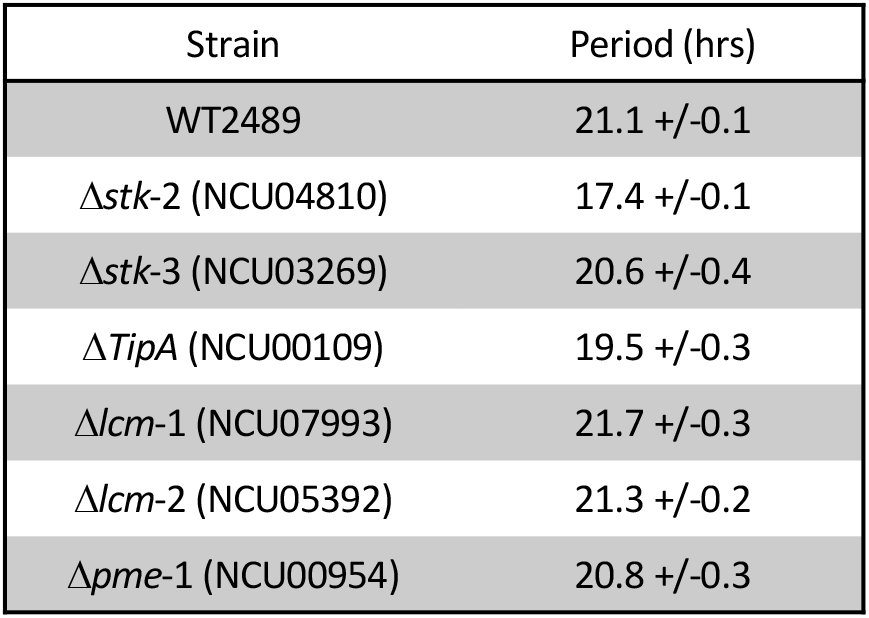
Period lengths of mutants in genes encoding PP4 regulatory proteins at 25°C. Known PP4 regulatory proteins in yeast and other fungal species were identified in Neurospora by homology; **see text for details**. Deletions were obtained from the Neurospora knockout collection at the FGSC and were crossed or transformed to insert the cbox-luc reporter at *csr*-1. Monitoring of luciferase activity demonstrates loss of *stk*-2 or *TipA* shortens the period. Loss of *stk*-3, or the leucine methyl-transferases / methyl-esterase acting on the PP4 C-terminal tail do not alter period.

### Mass spectroscopy suggests PP4 targets and uncovers a possible connection to protein chaperones

Immunoprecipitation of proteins followed by mass spectroscopy of the immunoprecipitate is commonly used to identify interacting proteins; we adapted this approach to identify potential substrates of the PP4 holoenzyme using the N-terminal VHF-tagged REG3 subunit for co-IP followed by LC-MS/MS. As the period of Δ*pp*4 changes with temperature, catalytic activity or substrate selection may also be temperature-dependent so lysates were prepared from cultures grown at 21°C and 29°C, along with untagged controls. A limited number of REG3 interactors were identified (Table 2) as has been seen before when this approach was used to identify PPP phosphatase interactors [37], and no interactions appeared to be altered by temperature. Proteins were identified by peptides that were over-represented as compared to controls, where the total peptide counts from each LC-MS/MS run are listed in Table 2. After excluding essential genes and those without available knockout strains, we obtained a final list of 12 interacting candidates; notably absent from this list are any known clock genes. Comparing this list to a PP4 specific LC-MS/MS experiment performed with cultured mammalian cells [25], and to the MCLR inhibitor LC-MS/MS experiment targeting all PPP family members which used *Neurospora* lysates [31], all of our candidates were novel candidate interacting proteins. Candidate interactors were screened by luciferase reporter assay to measure period at 25°C. All produced periods similar to wildtype controls, with the exception of the NCU07414 deletion. Loss of NCU07414 results in a stunningly short period of ~12-14 hours (Fig 5A), considerably shorter than the shortest *pp*4 knockout period under any conditions tested and as short or shorter than any allele of any gene previously examined in a circadian context in Neurospora.

**Table 2:** REG3 interacting proteins identified by LC-MS/MS. Immunoprecipitates of REG3 and interacting proteins were run on LC-MS/MS. Due to processing limitations, samples were divided between two preparations and therefore could not be combined for statistical tests. Instead, the counts of peptides corresponding to each protein are presented for the tagged REG3 strain and the wildtype control; peptide counts from independent experiments are separated by commas. Genes shown are those which are over-represented in the tagged strain. Deletion mutants for each gene were screened for circadian period using the cbox-luc reporter assay. All strains had periods similar to wildtype controls, with the exception of the NCU07414 knockout.

| N. crassa gene | Uniprot accession | Gene description (symbol) | @ 21C - n=4 per geno |  | @ 29C - n=3-4 per geno |  | KO period @25C (hrs) |
| --- | --- | --- | --- | --- | --- | --- | --- |
|  |  |  | Count of IP peptides | Count of peptides in untagged control | Count of IP peptides | Count of peptides in untagged control |  |
| NCU06389 | V5INY8 | DUF625 domain-containing protein | 70,61,104,102 | 0,0,0,0 | 52,77,95 | 0,0,0,0 |  |
| NCU08301 | Q7RVY0 | calcineurin-like phosphoesterase (*pp4*) | 21,21,26,26 | 0,0,0,0 | 18,23,21 | 0,0,0,0 |  |
| NCU00900 | Q7SCV4 | hypothetical protein | 40,42,47,58 | 0,0,0,0 | 34,42,35 | 0,0,0,0 |  |
| NCU00109 | U9W318 | TipA protein | 4,4,0,0 | 0,0,0,0 | 5,0,0 | 0,0,0,0 |  |
| NCU01953 | Q7SDY4 | AMI protein (*eat-3*) | 0,0,49,62 | 0,0,4,0 | 0,31,40 | 0,0,0,0 | 21.4 |
| NCU01747 | Q1K5R7 | glycerophosphocholine phosphodiesterase-1 (*gde-1*) | 0,0,11,13 | 0,0,0,0 | 1,14,19 | 0,0,0,0 | 21.9 |
| NCU00420 | U9WH02 | hypothetical protein | 0,0,6,6 | 0,0,0,0 | 0,4,7 | 0,0,0,0 | 20.7 |
| NCU07414 | Q7S1F9 | DnaJ family protein (*djc-4*) | 2,1,1,3 | 0,0,0,0 | 3,3,3 | 3,4,1,0 | ~12 |
| NCU05160 | Q7S8H3 | metalloprotease-15 (*mpr-15*) | 0,4,9,7 | 0,0,0,2 | 1,12,14 | 0,1,0,0 | 21.1 |
| NCU04047 | Q7RZG9 | hypothetical protein (*nup-19*) | 1,3,6,7 | 0,0,0,2 | 1,4,7 | 0,2,0,0 | 21.1 |
| NCU07574 | Q7S118 | hypothetical protein | 0,0,12,16 | 0,0,0,0 | 0,17,21 | 2,0,1,0 | 22 |
| NCU00695 | Q7SHV9 | hypothetical protein | 0,0,7,8 | 0,0,0,0 | 0,8,9 | 1,0,3,2 | 21 |
| NCU03725 | Q9C2N1 | VIB-1 (*vib-1*) | 0,0,4,4 | 0,0,0,0 | 0,1,2 | 0,0,0,0 | 20.5 |
| NCU04272 | V5IMT4 | zinc finger transcription factor-55 (*znf-55*) | 0,0,19,19 | 0,0,0,0 | 0,14,14 | 0,0,0,0 | 21 |

**Figure 5.**
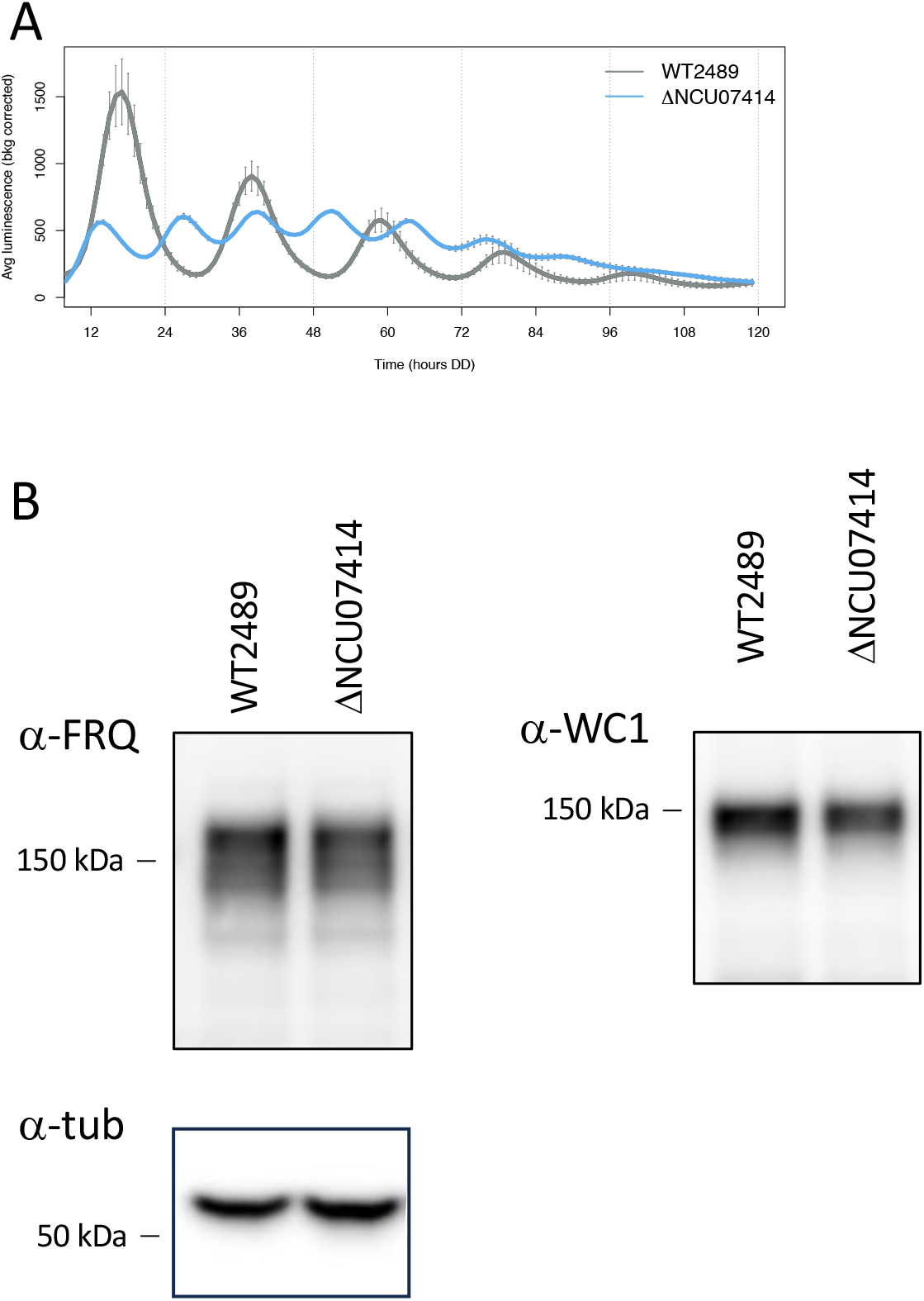
*djc*-4 (NCU07414) deletion results in an extremely short period of ~12 hrs. (A) Luciferase reporter assay of a *djc*-4 deletion strain at 25°C shows one of the shortest circadian periods known. (B) Abundance of FRQ and WC-1 are not appreciably altered in the *djc*-4 deletion. Strains were grown in 2% LCM under 25°C LL conditions. Western blots were probed with anti-FRQ, anti-WC-1 or anti-tubulin antibodies.

NCU07414 is annotated as DnaJ/Hsp40 family protein *djc-*4 and it is indeed similar with 66% identity to yeast Ydj1. Hsp40s are responsible for stabilizing interactions of Hsp70 chaperones with their client proteins and stimulating the chaperone folding ability. Homokaryotic deletions of NCU07414 were difficult to isolate from germinated ascospores as they grew more slowly than wildtype progeny at 30°C and were sensitive to presence of antibiotic selection despite the presence of antibiotic resistance genes. Biochemical analysis showed by Western blot that the abundance of core clock proteins WC-1 and FRQ was not altered in ΔNCU07414 (Fig. 5B). Careful analysis of prior LC-MSMS results [8] revealed a time-of-day-specific interaction between FRQ and DJC-4 peaking after 12 and 16 hrs in darkness; however, we were unable to confirm interactions with either FRQ, WC-1, or WC-2 in immunoprecipitations using an N-terminally VHF-tagged DJC-4 (data not shown). Plainly, loss of DJC-4 has a major impact on the clock but it seems unlikely that the effect is through direct interactions with known core clock components.

## DISCUSSION

We have identified the clock-relevant PP4 regulatory subunits in *Neurospora* and demonstrate that a heterotrimer of PP4/REG2/REG3 is required for phosphatase function. In the absence of PP4 activity clock rhythmicity is altered in a manner dependent on nutrition, where increasingly rich media shortens the period. PP4 is clearly a member of the unusual group of temperature and nutritional compensation mutants, at least in *Neurospora*. Only a few compensation mutants are identified [11, 38–42], and it remains to be seen if PP4 acts in concert with these or via a new mechanism.

We demonstrate the most likely site of action for PP4 is within the nucleus, based on in vivo localization of the REG3 subunit which binds target substrates of the phosphatase. In addition, we show that WCC localization remains predominantly nuclear, despite PP4 loss. This is contrary to a previous model of PP4 dephosphorylating WCC in the cytosol to facilitate nuclear entry of the transcription factor [17].

We identify a binding partner of DJC-4, which dramatically shortens the clock period to ~12hrs when deleted. DJC-4 is a HSP40 chaperone protein, which is itself regulated by phosphorylation. There are several possibilities for DJC-4 function in the clock: DJC-4 could be chaperoning the folding of PP4 and/or PP4 regulatory subunits. This seems unlikely given that period of the *djc*-4 deletion is even shorter than PP4 deletion. DJC-4 could be chaperoning the folding of core clock proteins, especially FRQ which is highly unstructured. In this case PP4 would be controlling the chaperone activity of DJC-4 through phosphorylation events. DJC-4 could be acting as a scaffold to associate PP4 phosphatase with the clock-relevant substrate. In this case ablation of the scaffold would decrease PP4 activity, similar to the effect of PP4 deletion.

That the DJC-4 deletion period is even shorter than that seen in a PP4 deletion hints this chaperone may have roles through other phosphatases. The PPP family of phosphatases, of which PP2A and PP4 are members, is highly conserved across evolution. Because of this similarity, interacting regulatory proteins are shared between some family members. It remains to be seen if DJC-4 interacts only with PP4 or if interactions with PP2A and/or PP6 are important for determination of circadian period and temperature compensation. Alternatively, it may be that PP4 helps in DJC-4 function or substrate recognition such that loss of PP4 represents a partial loss of DJC-4 activity and a short period whereas deletion of DJC-4 represents a complete loss and a more severe period shortening. In either case, period shortening is relatively unusual among clock mutant alleles and must represent loss of a normal delay in the feedback loop. In this case the delay would be instigated by phosphatase action such that the target phosphorylations would be acting to speed to clock rather than to slow its pace as is more typically the case for clock-relevant phosphorylations.

## MATERIALS AND METHODS

Neurospora strains were constructed using standard crossing and transformation methods. For insertion of tags, transformation cassettes were constructed using Gibson assembly for scarless sequence modification and amplified with a high-fidelity polymerase. Period of the core clock was measured using the cbox-luc reporter assay as previously described, with minor modifications to the growth media. Unless stated, solid agar growth media contained 0.01M QA. For glucose and QA titrations, water was substituted to replace the QA volume as necessary. Western blots were performed using 4-12% Bis-Tris gels with MOPS running buffer. Custom anti-FRQ and anti-WC1 antibodies were used as previously described. Anti-V5 antibody was purchased from BioRad. Anti-tubulin antibody was purchased from Sigma.

## ACKNOWLEDGEMENTS

We thank Dunlap-Loros lab members for helpful discussions. This work was supported by NIH R35GM118021 to JCD and R35GM145596 to SAG. The authors declare no conflicts of interest.

